# Serum CSF1 levels are elevated in patients following partial hepatectomy and acetaminophen induced acute liver failure: Reanalysis of previous data

**DOI:** 10.1101/2021.04.02.437789

**Authors:** BM Stutchfield, PJ Starkey Lewis, SJ Wigmore, K Simpson, DA Hume, SJ Forbes

## Abstract

A previous study analysed serum CSF1 concentration in patients who had undergone partial hepatectomy and patients with acute liver failure (ALF) induced by acetaminophen. A correlation between low CSF1 and poor prognosis was reported in ALF. In May 2018, an allegation of research misconduct was made against the second author of the published study, D.J.Antoine, who carried out the CSF1 analysis. The current study describes the independent reanalysis of circulating CSF1 concentration in sera from the same patient cohorts. Patient serum CSF1 levels increased following partial hepatectomy. By contrast to the published report, serum CSF1 concentration was greatest in ALF patients who died compared to survivors. The results confirm the relationship between liver injury and circulating serum CSF1.

## Introduction

Macrophage colony-stimulating factor (CSF1) is the main physiological regulator of proliferation, differentiation and survival of monocytes and tissue macrophages (1). Administration of recombinant human CSF1 to mice caused a monocytosis and expansion of tissue macrophage populations (2). Translation of this finding into therapeutic applications was constrained by the short half-life of the recombinant protein. This constraint was addressed through the development of an active CSF1-Fc fusion protein (3). CSF1-Fc administration to mice caused a rapid expansion of the size of the liver due to indirect effects of the expanded macrophage population on hepatocyte proliferation (3). In a subsequent study (4) we explored the role of CSF1-dependent macrophages in recovery from hepatic injury. Inhibition of CSF1 receptor (CSF1R) signalling impaired liver regeneration following partial hepatectomy, whereas CSF1-Fc treatment accelerated repair in both partial hepatectomy and acute injury models.

CSF1 in the circulation is rapidly cleared by CSF1R-mediated endocytosis by the macrophages of the liver and spleen (1). In the same published study (4), we reported on the detection of CSF1 in the circulation of patients undergoing partial hepatectomy or following admission with acute liver failure. This analysis suggested that circulating CSF1 had value as a prognostic indicator. In May 2018, we were made aware of an allegation of research misconduct against the second author of this paper, DJ Antoine (DJA) who conducted the biomarker work at a separate centre (University of Liverpool). An investigation was undertaken by the University of Liverpool commencing in Autumn 2017. The panel produced its final report in March 2018, which was published in July 2018 concluding that DJA was involved with research misconduct (https://news.liverpool.ac.uk/2018/07/06/research-misconduct-update/). In view of this author’s involvement in the analysis of the clinical material for our published study, we decided to re-analyse the samples. We therefore analysed patient samples from the cohorts included in the CSF1 paper which had been stored at the University of Edinburgh and DJA did not have access to. We report here that CSF1 is indeed elevated in the serum of patients following partial hepatectomy and in acute liver failure. However, the relationship to prognosis was the direct opposite to the previous conclusion; high CSF1 was associated with poor outcome.

## Methods

Serum from patients following partial hepatectomy included in the published CSF1 study (4) had been retained at the University of Edinburgh and stored at -80 degrees. Available serum from a cohort of patients with acetaminophen overdose who were included in the previous analysis but were retained separately at the University of Edinburgh was also analysed. A researcher independent to the prior CSF1 work (PS, Centre for Regenerative Medicine, University of Edinburgh) conducted the biomarker work using the human CSF1 ultrasensitive kit on the MSD platform. The independent reanalysis was undertaken blinded to the clinical history, grouping or outcomes of the patients. Results were analysed using Graphpad Prism v6. Two-tailed Student t test and 1-way and 2-way analysis of variance with Bonferroni adjustment were used for analysis of data. The level of significance was set at a P value less than .05 for all analyses.

## Results

Following partial hepatectomy in patients the previous analysis showed a reduction in serum CSF1 between preop and day 1 post op with a subsequent increase at day 3 post op. The reanalysis showed a progressive increase serum CSF1 levels from preoperative to day 1 postoperative and day 3 postoperative (Figure 1A). Consistent with published reports (5, 6), the serum CSF1 concentration detected was in the range of 10-100pg/ml, whereas the previous analysis of the same samples (4) reported concentrations in the ng/ml range. The previous analysis indicated a positive correlation between the number of segments resected and circulating CSF1 whereas reanalysis indicated the reverse relationship (Figure 1B).

**Figure 1.**
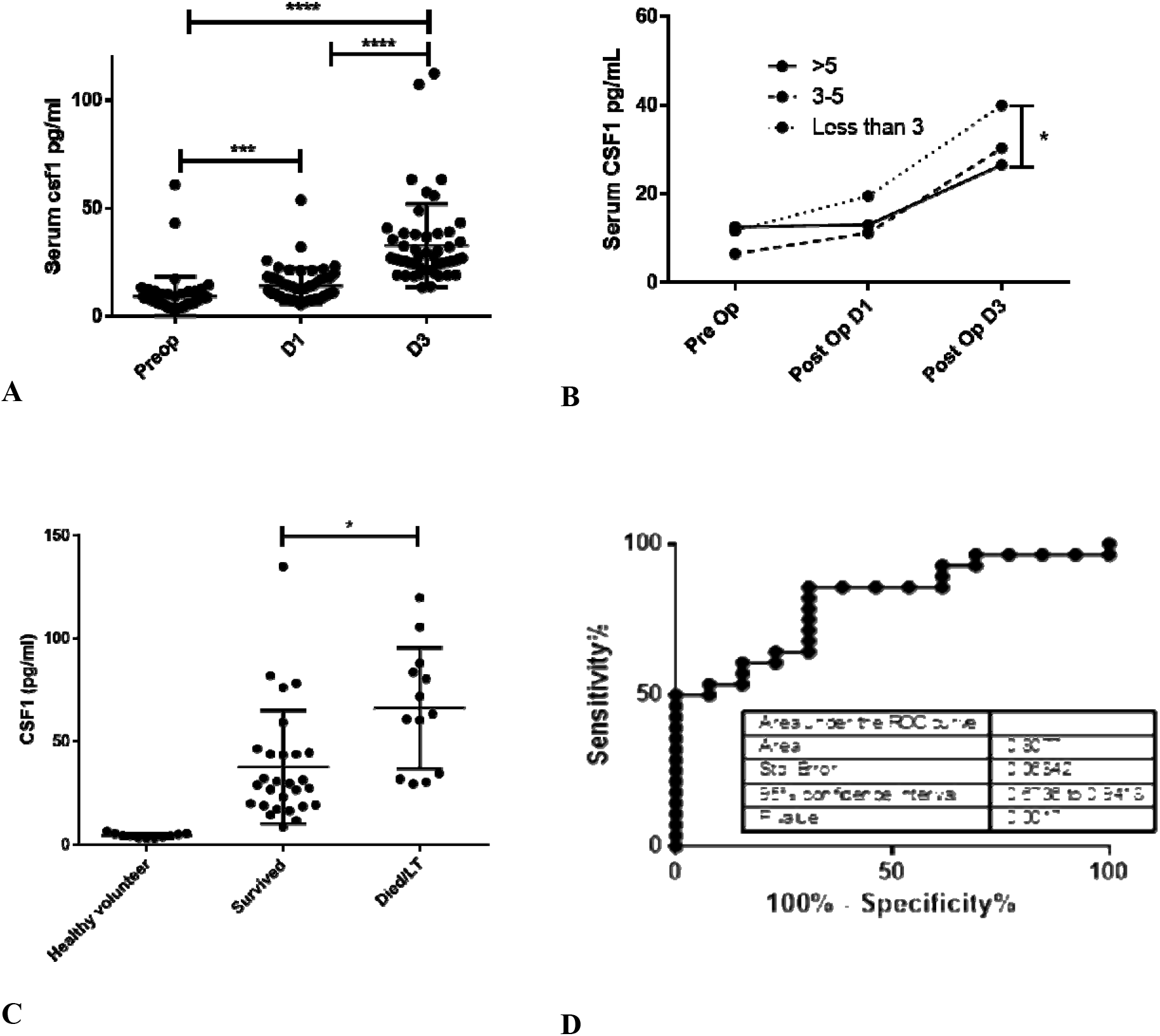
The effect of liver injury on circulating levels of CSF1. **A)** Serum CSF1 levels following partial hepatectomy (n=52) for all patients preoperatively, post-operative day 1 and post-operative day 3. **B)** Serum CSF1 for patients undergoing partial hepatectomy according to number of liver segments resected (more than 5 segments, 3-5 segments or less than 3 segments). **C)** Serum CSF1 analysis levels in healthy volunteers (n=11) and following presentation with acetaminophen intoxication (survived n= 29; died or liver transplant n=13) **D)** Receiver Operator Characteristic curve for serum CSF1 level following acetaminophen intoxication (AUC= 0.81, p=0.002)

Serum CSF1 level was reported to be highest in patients who survived acetaminophen overdose and was therefore proposed prognostic value (4). Reanalysis of the same samples revealed that the patients who died or underwent liver transplantation actually had higher CSF1 levels (Figure 1C). The ROC analysis showed area under the curve for the published analysis of 0.86. Based upon the new data, there was a significant correlation of serum CSF1 concentration with poor outcome and the ROC was 0.81 (Figure 1D).

## Discussion

Reanalysis of the CSF1 concentration in the serum samples of patients undergoing partial hepatectomy or in acute liver failure effectively reverses the previous conclusions about the relationship to liver injury severity and prognosis. Since there has been no explicit investigation of the generation of the previous data, we cannot conclude that these data were fraudulent. However, the range of detection reported (5-10 ng/ml) is well outside the range detected in other studies (5,6) including ongoing studies in patients with chronic liver disease (DAH, unpublished). The conclusion from reanalysis is consistent with a major role for hepatic macrophages in control of circulating CSF1 (1) and indicates that high CSF1 is an indication of the severity of liver injury. In that respect, CSF1 provides similar prognostic information to detection of HMGB1 (7). Others have reported elevated circulating CSF1 in chronic liver injury (5, 6).

The remaining findings in our published study using mouse models, indicating the therapeutic value of CSF1-Fc treatment for both recovery of innate immune function and hepatocyte regeneration, were conducted independently of any involvement by DJA and remain entirely valid. The ability of the pig CSF1-Fc protein used in the published study to promote hepatocyte proliferation has since been confirmed in rats (8) and pigs (9) and treatment was well-tolerated. Furthermore, both groups involved in the earlier study (DAH, BS/SF) have confirmed the efficacy of a GMP-grade mouse Fc-human CSF1 fusion protein in a range of liver injury models (Ms in preparation). We suggest that the elevated CSF1 seen in patients is a pro-repair response and CSF1-Fc has potential as a therapy in liver disease.

## References

1. Hume, D. A., M. Caruso, M. Ferrari-Cestari, K. M. Summers, C. Pridans, and K. M. Irvine. 2020. Phenotypic impacts of CSF1R deficiencies in humans and model organisms. J Leukoc Biol 107: 205–219.

2. Hume, D. A., R. E. Donahue, and I. J. Fidler. 1989. The therapeutic effect of human recombinant macrophage colony stimulating factor (CSF-1) in experimental murine metastatic melanoma. Lymphokine Res 8: 69–77.

3. Gow, D. J., K. A. Sauter, C. Pridans, L. Moffat, A. Sehgal, B. M. Stutchfield, S. Raza, P. M. Beard, Y. T. Tsai, G. Bainbridge, P. L. Boner, G. Fici, D. Garcia-Tapia, R. A. Martin, T. Oliphant, J. A. Shelly, R. Tiwari, T. L. Wilson, L. B. Smith, N. A. Mabbott, and D. A. Hume. 2014. Characterisation of a novel Fc conjugate of macrophage colony-stimulating factor. Mol Ther 22: 1580–1592.

4. Stutchfield, B. M., D. J. Antoine, A. C. Mackinnon, D. J. Gow, C. C. Bain, C. A. Hawley, M. J. Hughes, B. Francis, D. Wojtacha, T. Y. Man, J. W. Dear, L. R. Devey, A. M. Mowat, J. W. Pollard, B. K. Park, S. J. Jenkins, K. J. Simpson, D. A. Hume, S. J. Wigmore, and S. J. Forbes. 2015. CSF1 Restores Innate Immunity After Liver Injury in Mice and Serum Levels Indicate Outcomes of Patients With Acute Liver Failure. Gastroenterology 149: 1896–1909 e1814.

5. Preisser, L., C. Miot, H. Le Guillou-Guillemette, E. Beaumont, E. D. Foucher, E. Garo, S. Blanchard, I. Fremaux, A. Croue, I. Fouchard, F. Lunel-Fabiani, J. Boursier, P. Roingeard, P. Cales, Y. Delneste, and P. Jeannin. 2014. IL-34 and macrophage colony-stimulating factor are overexpressed in hepatitis C virus fibrosis and induce profibrotic macrophages that promote collagen synthesis by hepatic stellate cells. Hepatology 60: 1879–1890.

6. Shoji, H., S. Yoshio, Y. Mano, E. Kumagai, M. Sugiyama, M. Korenaga, T. Arai, N. Itokawa, M. Atsukawa, H. Aikata, H. Hyogo, K. Chayama, T. Ohashi, K. Ito, M. Yoneda, Y. Nozaki, T. Kawaguchi, T. Torimura, M. Abe, Y. Hiasa, M. Fukai, T. Kamiyama, A. Taketomi, M. Mizokami, and T. Kanto. 2016. Interleukin-34 as a fibroblast-derived marker of liver fibrosis in patients with non-alcoholic fatty liver disease. Sci Rep 6: 28814.

7. Gaskell, H., X. Ge, and N. Nieto. 2018. High-Mobility Group Box-1 and Liver Disease. Hepatol Commun 2: 1005–1020.

8. Irvine, K. M., M. Caruso, M. F. Cestari, G. M. Davis, S. Keshvari, A. Sehgal, C. Pridans, and D. A. Hume. 2020. Analysis of the impact of CSF-1 administration in adult rats using a novel Csf1r-mApple reporter gene. J Leukoc Biol 107: 221–235.

9. Sauter, K. A., L. A. Waddell, Z. M. Lisowski, R. Young, L. Lefevre, G. M. Davis, S. M. Clohisey, M. McCulloch, E. Magowan, N. A. Mabbott, K. M. Summers, and D. A. Hume. 2016. Macrophage colony-stimulating factor (CSF1) controls monocyte production and maturation and the steady-state size of the liver in pigs. Am J Physiol Gastrointest Liver Physiol 311: G533–547.

